# GENERATION AND CHARACTERIZATION OF IMMORTALIZED MOUSE CORTICAL ASTROCYTES FROM WILDTYPE AND CONNEXIN43 KNOCKOUT MICE

**DOI:** 10.1101/2020.12.28.424504

**Authors:** Antonio Cibelli, Sandra Veronica Lopez-Quintero, Sean McCutcheon, Eliana Scemes, David C. Spray, Randy F. Stout, Sylvia O. Suadicani, Mia M. Thi, Marcia Urban-Maldonado

**Author notes:** Department of Mechanical Engineering, Manhattan College, Riverdale, NY. Department of Anatomy and Cell Biology, New York Medical College, Sunshine Cottage Rd, Valhalla, NY. **Corresponding Author:** Dr. Randy F. Stout Jr, Department of Biomedical Science, New York Institute of Technology College of Osteopathic Medicine’, New Westbury, NY.

## Abstract

We transduced mouse cortical astrocytes cultured from four litters of embryonic wildtype (WT) and connexin43 (Cx43) null mouse pups with lentiviral vector encoding hTERT and measured expression of astrocyte-specific markers up to passage 10 (p10). The immortalized cell lines thus generated (designated IWCA and IKOCA, respectively) expressed biomarkers consistent with those of neonatal astrocytes, including Cx43 from wildtype but not from Cx43-null mice, lack of Cx30, and presence of Cx26. AQP4, the water channel that is found in high abundance in astrocyte end-feet, was expressed at moderately high levels in early passages, and its mRNA and protein declined to low but still detectable levels by p10. The mRNA levels of the astrocyte biomarkers aldehyde dehydrogenase 1 (ALDH1), glutamine synthetase (GS) and glial fibrillary acidic protein (GFAP) remained relatively constant during successive passages. GS protein expression was maintained while GFAP declined with cell passaging but was still detectable at p10. Both mRNA and protein levels of glutamate transporter 1 (GLT-1) declined with passage number. Immunostaining at corresponding times was consistent with the data from Western blots and provided evidence that these proteins were expressed at appropriate intracellular locations. Consistent with our goal of generating immortalized cell lines in which Cx43 was either functionally expressed or absent, IWCA cells were found to be well coupled with respect to intercellular dye transfer and similar to primary astrocyte cultures in terms of time course of junction formation, electrical coupling strength and voltage sensitivity. Moreover, barrier function was enhanced in co-culture of the IWCA cell line with bEnd.3 microvascular endothelial cells. In addition, immunostaining revealed oblate endogenous Cx43 gap junction plaques in IWCA that were similar in appearance to those plaques obtained following transfection of IKOCA cells with fluorescent protein tagged Cx43. Re-expression of Cx43 in IKOCA cells allows experimental manipulation of connexins and live imaging of interactions between connexins and other proteins. We conclude that properties of these cell lines resemble those of primary cultured astrocytes, and they may provide useful tools in functional studies by facilitating genetic and pharmacological manipulations in the context of an astrocyte-appropriate cellular environment.

## Introduction

Vertebrate gap junctions (GJs) are formed by the connexin family of proteins. Gap junction channels provide direct cytosolic exchange of current-carrying ions, metabolic substrates and signaling molecules between adjacent cells. The most widespread and abundant gap junction protein is connexin43 (Cx43), which is the primary gap junction protein between cardiac myocytes, vasculature and astrocytes in the brain. Intercellular communication through Cx43 GJs mediates key essential functions in the tissues in which it is expressed, including propagation of contraction throughout the heart, signaling during migration of neural precursor cells, and coordination of cellular functions in communication compartments. Because it is such an abundant GJ protein, it was not surprising that when the first Cx43-null mouse was generated, the homozygous null pups died soon after birth(1). What was surprising was that the perinatal mortality was not the result of disturbed rhythmic propagation throughout the heart, but instead was due to delayed neural crest cell migration and ensuing developmental defect of the left ventricular outflow tract (2). As a consequence, the *foramen ovale* between the left and right cardiac chambers in Cx43-null mice does not close at birth. Thus, Cx43-null mice survive until placental circulation ends, at which time the animals are no longer adequately perfused.

In order to study the properties of cells lacking Cx43 expression, we maintained breeding colonies of Cx43 heterozygotes for many years so that we could obtain tissues and prepare primary cell cultures from wildtype and Cx43-null P0 pups immediately after delivery, or from E19-20 pups obtained by c-section (3). This protocol has been costly and inefficient, as breeding is irregular in these transgenic mice, and cell cultures must be separately prepared from each individual mouse before genotype is verified. In order to circumvent these difficulties, we have transduced primary cultures of cortical astrocytes from Cx43-null and wildtype littermates with human telomerase reverse transcriptase (hTERT). This method has been successfully used to generate other cell lines, including wildtype and Cx43-null osteoblastic cell lines, as we previously described (4). In addition, hTERT transduction is reported to have the advantage over transfection with oncogenes in that it does not involve the inactivation of tumor suppressor genes (5).

In this manuscript, we report the characterization of hTERT-immortalized astrocyte cell lines derived from wildtype and Cx43-null mice, designated IWCA and IKOCA cells, with respect to expression of astrocyte-specific mRNAs and proteins as well as several diverse functional features. We also demonstrate that GJ plaques formed of Cx43 exogenously expressed in IKOCA cells are similar in appearance to GJ plaques expressed in the wildtype primary astrocyte cell culture, indicating the utility of these cell lines for studies of the impact of cellular and molecular manipulations on astrocyte structure and function.

## Material and Methods

### Cell cultures

We used primary cultures of cortical astrocytes derived from 19-20 day old (E19-20) wildtype (WT) and Cx43-null mouse embryos (offspring of Cx43 heterozygotes in C57Bl/6J-Gja1 strain, originally obtained from Jackson Laboratories). Animals were maintained at the Albert Einstein College of Medicine animal facility and all experimental procedures were approved by the college’s Institute of Animal Care and Use Committee. Cortices were separated from whole brain E19-E20 embryos and, after removal of meninges, tissues were minced and enzymatically digested (0.05% trypsin at 37°C for 10 min). Cells from each animal were collected by centrifugation and pellets suspended in Astrocyte Medium [ScienCell Research Laboratory (SCRL), cat# 1801], supplemented with 2% fetal bovine serum (FBS) (SCRL cat# 0010), 1% penicillin/streptomycin (SCRL cat# 08030) and 1% astrocyte growth factor (SCRL cat# 1852), and seeded in plastic culture dishes. Genotypes of the astrocyte culture obtained from each mouse pup were determined by PCR on tail DNA as described(6). Astrocytes were maintained for 10-14 days in culture (100% humidity; 95% air / 5% CO_2_, 37°C) until confluency, at which time cells were immortalized as we previously described for osteoblasts (4).

### Immortalization

We used the human telomerase reverse transcriptase (hTERT) overexpression system for immortalization of astrocytes. hTERT cDNA was PCR amplified from its original construct hTERT-pGRN145 (ATCC, MBA-141; Manassas, VA) and subcloned into pLentiV5-EF1α vector (modified from original vector purchased from Invitrogen, cat# V49610). Confluent cultures of WT and Cx43-null astrocytes were transduced with lentiviral particles containing hTERT cDNA 9×10^5^ transduction units (TU; particles/ml). After overnight incubation, the mixture was replaced with Astrocyte Medium. Selection of immortalized cells was then achieved by successive splitting bi-monthly, thus eradicating cells that did not continue to divide. Cells that were successfully transduced with hTERT viral particles were designated as immortalized beginning at passage 2 because primary astrocytes grow only very poorly beyond second passage. The immortalization process was performed four independent times in different litters to generate four cell lines of immortalized WT cortical astrocytes (IWCA) and four cell lines of immortalized Cx43-null cortical astrocytes (IKOCA).

### Quantitative Polymerase Chain Reaction (qPCR)

The qPCR was performed using SYBR GREEN Master Mix (Applied Biosystems) in an Applied Biosystems 7300 Real-Time PCR System (Forester City, CA), according to the manufacturer’s instructions. Briefly, 2 μg total RNA was reverse transcribed into cDNA using SuperScript VILO cDNA Synthesis (Invitrogen). Amplification was carried out for 40 cycles with annealing temperature of 60°C in 25 μl final volume. Finally, a dissociation profile of the PCR product(s) was obtained by a temperature gradient running from 60°C to 95°C. The primers used for qPCR are listed in Table 1. Two technical replicas for each of the four immortalized WT and Cx43-null astrocyte cell lines were amplified. Relative gene expression levels of the analyzed mRNA samples were calculated using the ΔΔCt method, where values obtained for the gene of interest are first normalized to those of the reference, housekeeping gene, β-actin, and subsequently to those of their respective controls (non-hTERT-immortalized cells – passage 1). As in our previous studies(4), validation of the comparative Ct method was obtained after estimation of equality of efficiencies of target and reference amplifications (ΔCt (Ct_target_ – Ct_reference_)) over serial sample dilutions. Cells were retained from early passages and selected later passages as indicated for comparison of biomarker expression.

**Table 1:**
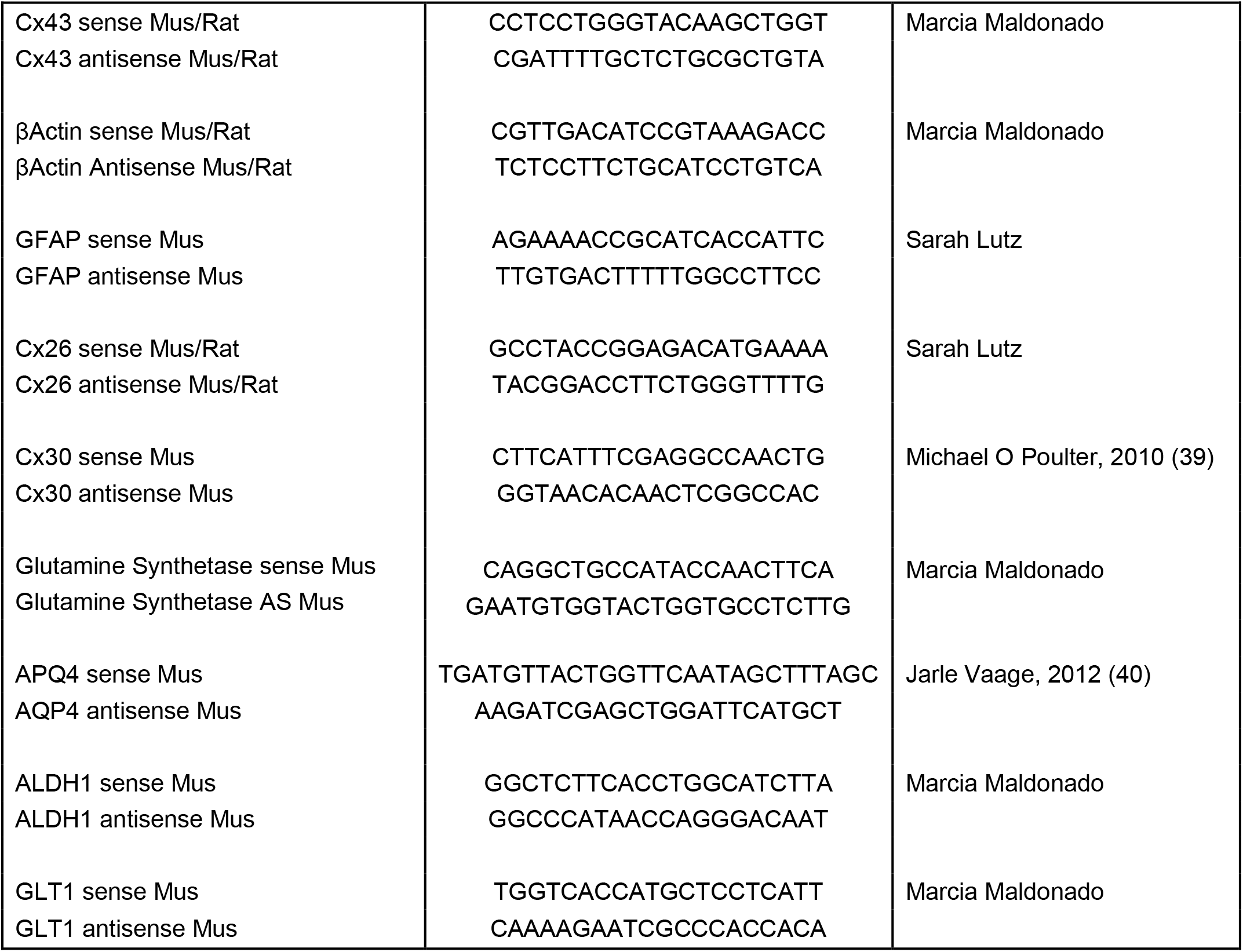
qPCR primer sequences used for characterization of immortalized astrocytes.

### Western blot analysis

Confluent astrocyte cultures plated in 100 mm dishes were lysed and equal amounts of total protein were loaded onto 7.5 or 10% SDS-PAGE gels for separation and electrophoretically transferred to nitrocellulose membranes (Whatman, Dassel, Germany). The membranes were probed with polyclonal antibodies to rabbit anti-Cx43, 1:10,000 (Sigma, cat# C6219); mouse anti-GFAP 1: 500 (Sigma cat# G3893), goat anti-GS 1:500 (Santa Cruz, cat# SC-6640), and goat anti-AQP4 1:500 (Santa Cruz, cat# SC-9888), followed by secondary antibody incubation with horseradish peroxidase-conjugated donkey anti-goat IgG, goat anti-rabbit IgG and goat anti-mouse (1:10,000, Santa Cruz Biotechnology, Santa Cruz, CA, cat# SC-2020, SC-2004 and SC-2005 respectively). Protein bands were detected using the Immobilon Western Chemiluminescent HRP Substrate (EMD Millipore, MA catalog# WBKLS0100) and images were acquired using the In Vivo FX PRO imaging system (Carestream, Carestream, NY, USA).

### Dye coupling

We used both dye injection of Lucifer Yellow (LY) and the so-called “parachute assay” with calcein-AM loaded donor cells to evaluate GJ-mediated intercellular diffusion of moderately large molecules (LY=444 Da; calcein=622 Da). Dye-injection was performed in astrocyte cultures of similar confluency. LY was iontophoresed (continuous current of 0.1 μA) for 5 min using an electrometer (model 3100; A-M Systems) into single cells using sharp microelectrodes filled with the dye (LY, 5% wt in 150 mM LiCl; resistance ~100 MOhms). After removal of the micropipette, images were immediately acquired using a CoolSNAP-HQ2 CCD camera (Photometrics, Tucson, AZ) attached to a Nikon inverted microscope with 10X dry objective (numerical aperture 0.3) and FITC filter sets. Fluorescent cells surrounding the LY-injected cell were counted, and the values were then normalized by the total number of cells within the injected region of interest (ROI) and expressed as percentage of coupled cells per ROI. The parachute assay was performed essentially as described (7, 8). In brief, cells in culture were detached via trypsinization (Thermo Fisher Scientific, #25200056), loaded in suspension with 10 μM Calcein-AM (Thermo Fisher Scientific, #C3100MP) and labeled with the lipophilic GJ impermeable dye DiI (Sigma Aldrich, #468495-100MG) for 30 min to distinguish the parachute donor cells from the recipient cells. Donor cells were then rinsed with Dulbecco’s phosphate buffer saline (DPBS, Mediatech, Cellgro, VA) and centrifuged twice to remove residual dye, then added to confluent monolayers of unlabeled recipient cells. Number of recipient cells adjacent to the donor cells stained with GJ permeant Calcein dye was determined for paired IWCA-IWCA, IKOCA-IKOCA and heterocellular IWCA-IKOCA. This assay is both confirmatory and supplemental to the LY injections, as it depends both on rate of GJ formation and on permeability of the GJs that are formed.

### Electrical coupling

The dual whole-cell voltage-clamp technique was used to characterize junctional conductance in IWCA and IKOCA cells, as described (9). Pairs of astrocytes were voltage-clamped at holding potentials of 0 mV, and 8–10 sec duration command steps (ΔV) in 20 mV increments from −110 to +110 mV or from −100 to +100 mV were presented to one cell with pClamp software (Axon Instruments, Foster City, CA). In some experiments a voltage ramp protocol was used, in which voltage slowly increased from −100 to +100 mV. Junctional currents (Ij) were recorded in the unstepped cell, and junctional conductance (Gj) was calculated as −Ij/ΔV (9). Patch pipettes were filled with (in mM) 140 CsCl, 10 EGTA, and 5 Mg_2_ATP, pH 7.25. Cells were bathed in solution containing (in mM) 140 NaCl, 2 KCl, 2 CaCl_2_, 1 BaCl_2_, 2 CsCl, 1 MgCl_2_, and 5 HEPES, pH 7.2.

### Transient exogenous expression of fluorescent protein-tagged Cx43

IKOCA cells were transfected with superfolderGFP-tagged Cx43 msfGFP-rCx43, EBFP2-rCx43, sfGFP-Cx30, sfGFP-Cx26 (10, 11) and msfGFP-ATG9A. This recovered Cx43 expression to allow comparison of exogenously expressed Cx43 aggregates to endogenously expressed plaques in primary WT astrocyte cultures. Cells were plated to 50-60% confluence in DMEM (Thermo Fisher Scientific, # 11885084) supplemented with 10% FBS and 1% penicillin/streptomycin (Thermo Fisher Scientific, # 15070063). Two hours prior to transfection, the media was changed. For transfections, 1 μg DNA was combined with 100 μL Opti-MEM (Thermo Fisher, #31985062) and 5μL transfection reagent (Mirus TransIT LT1, #MIR2300) and incubated at room temperature for 20-30 min. Liposome-DNA complexes were then added dropwise to IKOCA monolayers and incubated in standard culture conditions without changing media for 2-4 days before imaging.

### Confocal microscopy to analyze distributions of astrocyte proteins and to compare endogenous Cx43 junctions with those formed by fluorescent protein-tagged Cx43

To evaluate protein localization within the WT and Cx43-null primary astrocytes and immortalized cell lines, cultures plated on coverslips were fixed with 4% paraformaldehyde (PFA), permeabilized with 0.4% Triton-X100, and blocked with 2% donkey serum (Jackson Immunoresearch, West Grove, PA) as previously described(12). The cells were then incubated with primary antibody against rabbit anti-Cx43, 1:1,000 (Sigma, cat# C6219); mouse anti-GFAP 1: 500 (Sigma cat# G3893), goat anti-GS 1:500 (Santa Cruz, cat# SC-6640), and goat anti-AQP4 1:500 (Santa Cruz, cat# SC-9888), and secondary antibody conjugated to Alexa-488 goat-anti-mouse IgG (cat#A11001), Alexa Fluor 594 Donkey Anti-goat (cat#A-11058) and Alexa-488 goat anti-rabbit IgG (cat# A-11034) (1:1000, Invitrogen). The coverslips were mounted on slides, examined on a Zeiss LSM 510 DUO Laser Scanning Confocal Microscope (Carl Zeiss) with a 40x water-immersion objective. Stacked images were taken serially at 0.6 μm z axis steps.

Cx43 was either immunolabelled with primary rabbit polyclonal antibody (1:1000; Abcam, #ab11370) in permeabilized cells, as described above, or was identified by the fluorescent signal of GFP-tagged Cx43 (green fluorescent protein) in transfected cells. Image acquisition and z-stacks of immunolabeled junctional plaques were obtained with Zeiss LSM 880 inverted microscope with a 63x oil objective (numerical aperture, 1.4). Z-stacks of GJ plaques of exogenously expressed GFP-tagged Cx43 in live cells were obtained at 37°C with Zeiss 5Live Duo with 63x oil immersion objective. Imaging media for live cells was phenol-free DMEM (Thermo Fisher Scientific, # 31053028) supplemented with 10% FBS, 1% PS, and 25 mM HEPES (Thermo Fisher Scientific, # 15630080).

### Imaging of IKOCA cells transfected to express fluorescent protein-tagged connexins with co-staining via immunofluorescence

IKOCA cells were grown in Ibidi 8-well chambered ibiditreat chambered slides (Ibidi Inc,) and transfected with EBFP2-Cx43 then than 48 hours after transfection the were fixed in 4% PFA for 10 minutes, permeabilized with 0.4% Triton-X100, and blocked with 2% donkey serum and then immunostained using rabbit antibody against Cx43 (Figure 6A, Sigma, cat# C6219, 1:500) or Alpaca antibody against Green Fluorescent Protein conjugated to Alexa Fluor 488 (also labels EBFP2, Figure 6B, Chromotek, cat# gb2AF488, 1:100 dilution) to enhance the EBFP2 signal in the green color channel along with DAPI (blue to stain nuclei. To co-label mitochondria we stained with TOMM20 primary antibody (Invitrogen, cat# MA5-32148, 1:500) followed by secondary staining with Donkey anti-rabbit Alexa Fluor 647 (Figure 6B, Invitrogen, cat# A-31573, 1:500). We stained the cells in Figure 6A with Alexa 488 goat anti-rabbit IgG (Invitrogen, cat# A-11034, 1:500). Primary antibody labeling was performed for 1 hr at room temperature with agitation then overnight at 4 degrees. Secondary antibody staining was done for 3 hrs at room temperature with agitation. Images in Figure 6 were captured using a Zeiss 980 Airyscan2 laser scanning microscope with a 63x 1.4NA Plan-Apochromat oil immersion objective. The zoomed-out image in Figure 6B has a non-linear intensity lookup table for the green channel to allow the reader to simultaneously see both the very densely labeled reflexive GJ plaque and the non-junctional membrane localized Cx43. All other image lookup tables are scaled linearly by signal intensity.

### Live imaging and Fluorescent Recovery After Photobleach experiments on IKOCA cells expressing fluorescent protein-tagged proteins via transient transfection

IKOCA cells were grown in Ibidi 8-well chambered slides transfected as described for previous sections. sfGFP-Cx43, sfGFP-Cx30, sfGFP-Cx26 were used as described previously for other cell types (Stout et. al. 2015). Cells were transferred to imaging media containing 10% fetal bovine serum, 25 mm HEPES, and 2 mm glutamine in DMEM without phenol red. Live experiments in Figure 7 were carried out on the Zeiss 5live Duoscan 510 confocal laser scanning microscope with the 63x 1.4NA objective. 3D FRAP experiments were carried out by centering an 11 plane Z-stack acquisition on a gap junction plaque then acquiring an 11-plane 3D image every 3 seconds. A stripe photobleach was performed with the 488 nm laser at 100% with 3 iterations. The resulting 3D-timelapse image was converted to a maximum projection orthogonal reconstruction to produce a single-plane time-lapse image series in Figure 7A. For Figure 7B, IKOCA cells were co-transfected with EBFP2-Cx43 (McCutcheon et al 2020) along with human ATG9A tagged on the amino-terminus with monomeric superfolder GFP (msfGFP) produced by standard subcloning techniques, with plasmid sequences available on request. Cells were then imaged with 11 plane Z-stack acquisitions with both blue and green channels sequentially imaged per focal plane. The resulting two-color, 3D time-lapse was collapsed to a time-lapse sequence of maximum projection orthogonal projections for display in Figure 7.

### Measurement of barrier formation in IWCA-endothelial co-cultures

Transwell polyester inserts with 0.4 μm pores (Sigma Aldrich, # CLS3460) were flipped up-side down and coated with 100 μL of 30 μg/mL fibronectin (FN; Sigma Aldrich, # F1141) solution in DPBS. Coated inserts were incubated for 1 hour at 37°C in a humidified incubator with 95% air / 5% CO_2_. IWCA were then seeded at a density of 30,000 cells/cm^2^ and cells allowed to adhere to the FN coated Transwell insert for 2 hours, after which the inserts were flipped again and placed in 12 multi-well plates, with the IWCA cells now facing the bottom of the plate, and incubated for 2 days in DMEM. Mouse brain endothelial cells (bEnd.3 cells, ATCC CRL-2299, Manassas, VA) were then seeded on the luminal side of the insert at a density of 60,000 cells/cm^2^ and grown in DMEM. Companion inserts without IWCA were seeded with bEnd.3 alone to compare their barrier function to that of the co-cultures. Trans-endothelial electrical resistance (TEER) of endothelial monolayers and co-cultures of endothelial monolayers with IWCA was measured using the cellZscope system (Nanoanalytics, Munster, Germany) for automated, long-term monitoring of barrier resistance. Each insert seeded with cells was placed in a chamber filled with media (DMEM - 10% FBS), and the chamber holder was moved to the incubator (37°C, 5% CO_2_) and connected to the controller. CellZscope software measured TEER every 6 hours. Media in the chamber and the inserts were replaced every 3 to 4 days and TEER was monitored for up to 7 days after seeding the bEnd.3 cells on the insert.

### Statistical Analysis

GraphPad Prism 8 software was used for data and statistical analyses. Data are represented as mean ± SEM and considered statistically significant at *P<0.05. Statistical differences were determined by unpaired Student’s t-test for dye injection, parachute assay analyses, junctional conductance and day 7 of TEER. One-way ANOVA followed by Dunnett’s multiple comparison test was used for Western blot analysis.

## Results

### Quantification of expression levels of astrocyte markers in primary cultures of WT and Cx43-null astrocytes, and at the passages after each genotype was transfected with hTERT

Cx43 transcript expression was found to be relatively constant over time in IWCA astrocytes and was not at all detectable in IKOCA nulls (Fig.1A). Cx30 levels were too low to be quantified in both genotypes at all passages (Fig.1B), whereas Cx26 abundance was relatively constant in both genotypes up to passage 10 (Fig. 1C). AQP4 mRNA showed similarly sustained expression (Fig. 1D). Other astrocyte markers that we examined included glutamine synthetase (GS), aldehyde dehydrogenase 1 (ALDH1), glutamate transporter 1 [GLT-1, mouse ortholog of human excitatory amino acid transporter 2 (EAAT2) and solute carrier family 1 member 2 (SLC1A2)] and glial fibrillary acidic protein (GFAP). GS, ALDH1 and GFAP were found to have virtually constant mRNA expression levels over time in culture in both genotypes (Figs 1E, F, H). Expression of GLT-1 was present in two of the cell lines of both genotypes and below detection in the other 2 cell lines (Fig 1. G).

**Figure 1.**
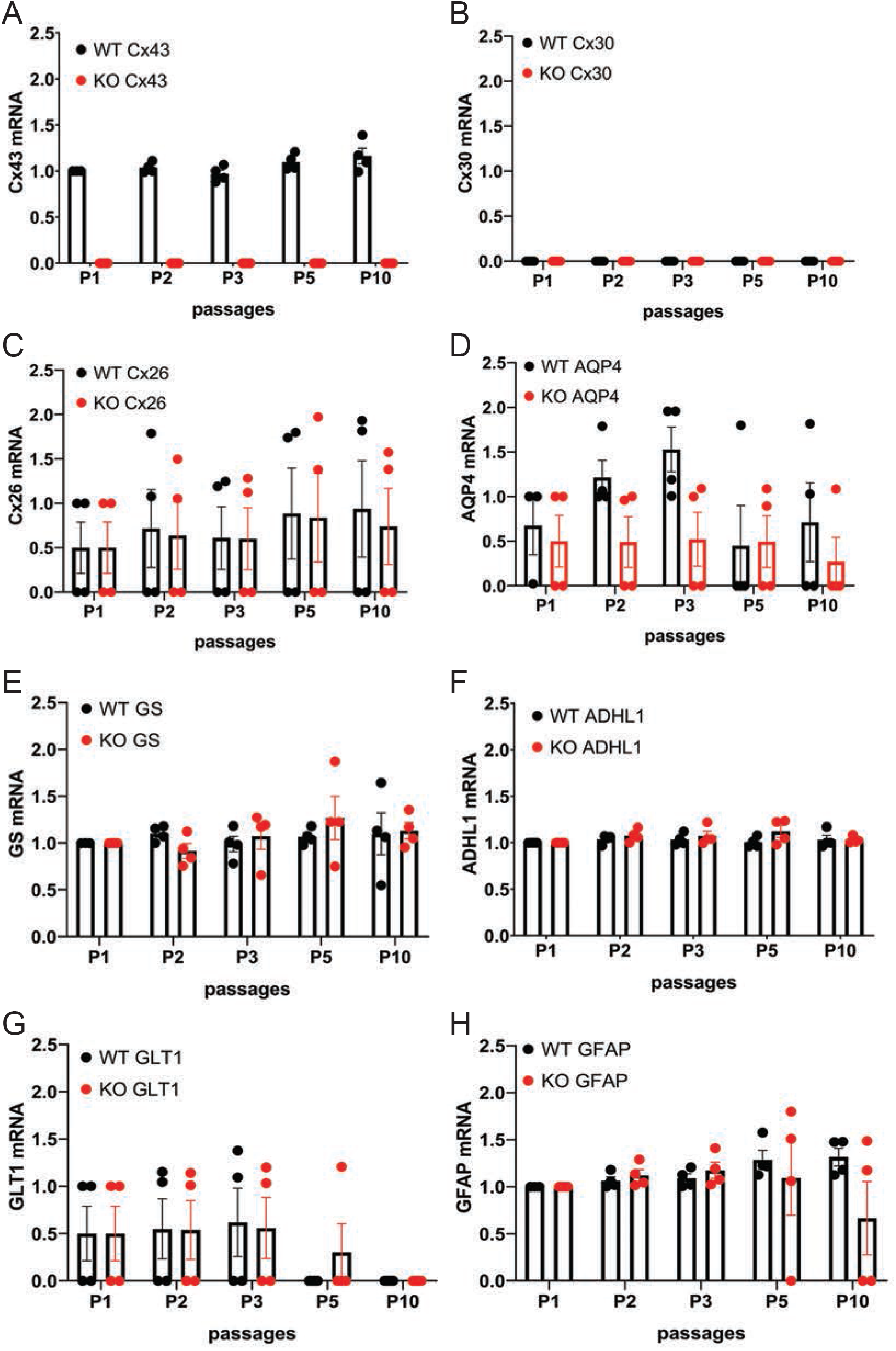
Quantitative RT-PCR showing expression levels of mRNAs encoding astrocyte proteins during ten passages after transduction. Expression levels of astrocyte biomarkers in the immortalized wildtype (IWCA) and Cx43-null (IKOCA) cortical astrocyte cell lines. Expression levels measured at each time point up to 10 passages after transduction with hTERT are normalized to those of IWCA astrocytes before transduction (labeled passage 1 in the histograms). Points in histograms correspond to individual cell lines and bars and variance correspond to mean ± SE values from the four independent clones generated for each of the two genotypes. **A**. Cx43 levels are stable in IWCA up to 10 passages, whereas Cx43 expression is absent in all of the IKOCA lines. **B**. Cx30 was not detectable in either IWCA or IKOCA, either before immortalization or at any time point thereafter. **C**. Cx26 is present in both IWCA and IKOCA and is relatively stable in both genotypes with continued passaging. **D**. Levels of AQP4 transcript in both IWCA and IKOCA, with the lowest levels at passage 10 being about 20% that of those in primary cells. **E and F**. Transcript levels of the astrocyte markers glutamine synthetase (GS) and aldehyde dehydrogenase (ALDH1) were remarkably stable over the 10 passages examined. **G.** mRNA for the glutamate transporter (GLT1) were lower than in primary cells but similar between genotypes and declined after passage 5 in both genotypes to reach undetectable levels at passage 10. **H**. Levels of the astrocyte intermediate filament transcript GFAP was stable in both genotypes but was low in two of the IKOCA cell lines at passage 10.

Western blots and immunostaining were performed to quantify protein expression levels and determine cellular localization in the IWCA and IKOCA at passages 1-10. Immunoblots and immunostaining for Cx43 at representative passages are shown in Figs. 2A, B and C, and for AQP4 in Figs. 2A, B and D. Cx43 protein level was not significantly reduced until passage 5 in the IWCA cultures but was reduced to about 30% of its initial level at passage 10 (Fig. 2C). Cx43 was not detected at any time point in immunostained cultures or in immunoblots obtained from IKOCAs. Both immunostaining and Western blots revealed that levels of AQP4 protein were lower in IKOCA than IWCA astrocytes at passage 1, and declined progressively in both genotypes from passage 1 onward (Fig. 2D).

**Figure 2.**
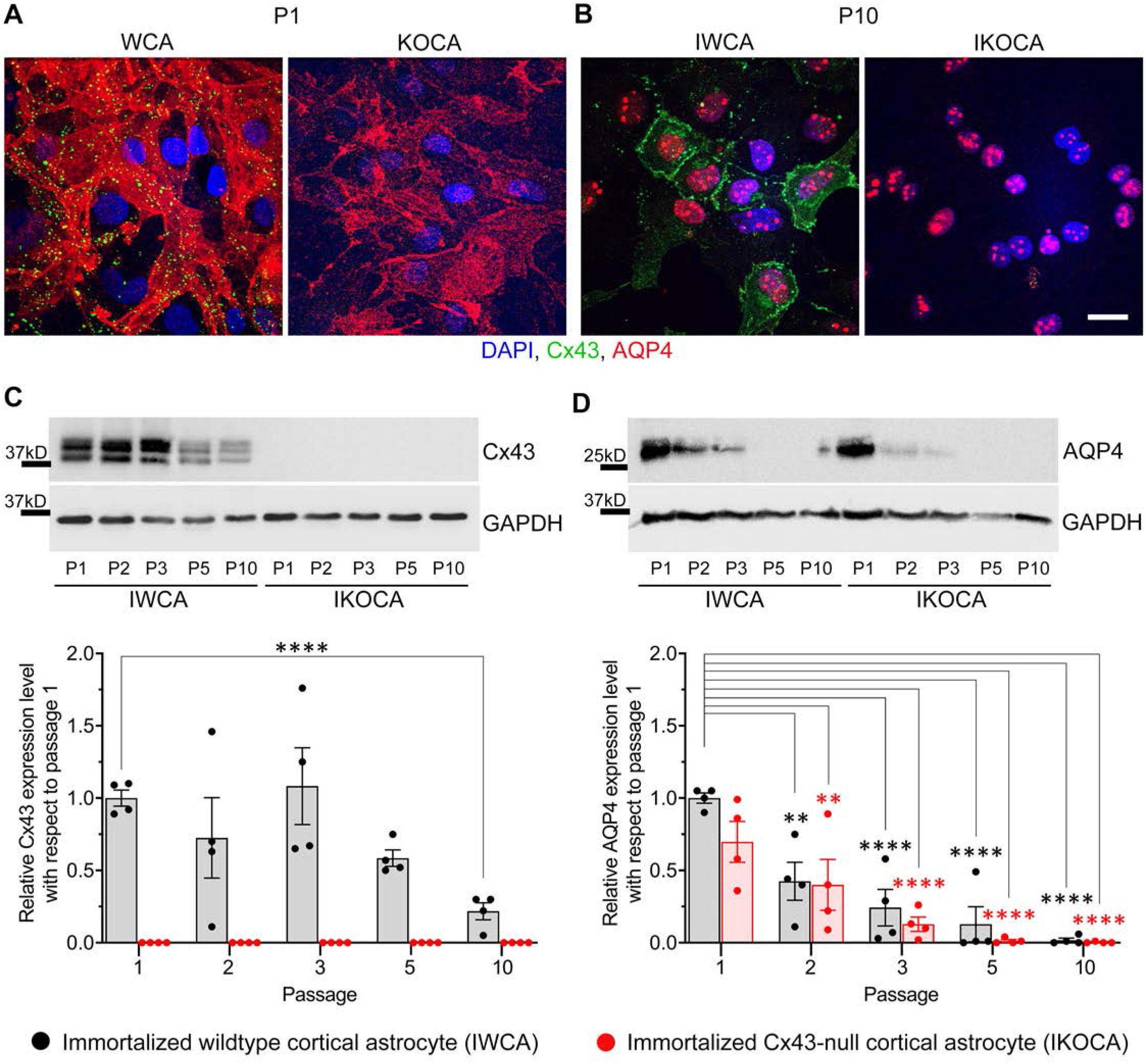
Expression level and localization of Cx43 and AQP4 in IWCA and IKOCA during the first ten passages. **A, B.** Immunostaining of Cx43 in immortalized WT (IWCA) and Cx43-null (IKOCA) cortical astrocyte cultures revealed punctate intercellular localization in IWCA that was less abundant but still prominent over time in culture. IKOCA showed no Cx43 immunostaining at any time point. Punctate distribution of AQP4 indicated that this protein was still well expressed at passage 10 in both IWCA and IKOCA, but at reduced to very low levels. Note: dot like staining in the nuclei staining is non-specific, likely as artifact of AQP4-C19 antibody (Ref. Thi et al., 2008 PMC2713861) **C**. Western blots showed no significant decline in Cx43 protein levels in IWCA until passage 10, when its level was reduced to about 30% of that of primary astrocytes. Cx43 protein expression was absent in IKOCA at all time points. **D.** AQP4 protein expression was very low but detectable at passage 5 and beyond in both genotypes.

Expression levels and intracellular staining patterns were also determined for GFAP and GS, two other characteristic astrocyte cell markers. Similar to mRNA results shown in Fig. 1, GFAP staining was detectable in both IWCA and IKOCA at all passages (Figs. 3A, B), but protein levels declined over time (Fig. 3C). However, this decline in GFAP protein expression was more rapid in the IKOCA than in IWCA, as revealed by the immunoblots (Fig. 3C). GS showed strong immunostaining in both IWCA and IKOCA until passage 10 (Figs. 3A, B). Western immunoblots detected an apparent increase in GS seen in both IWCA and IKOCA cultures until passage 5, which decreased thereafter (Fig. 3D). Variance was high, and differences between genotypes or over time were not significant as evaluated by ANOVA.

**Figure 3.**
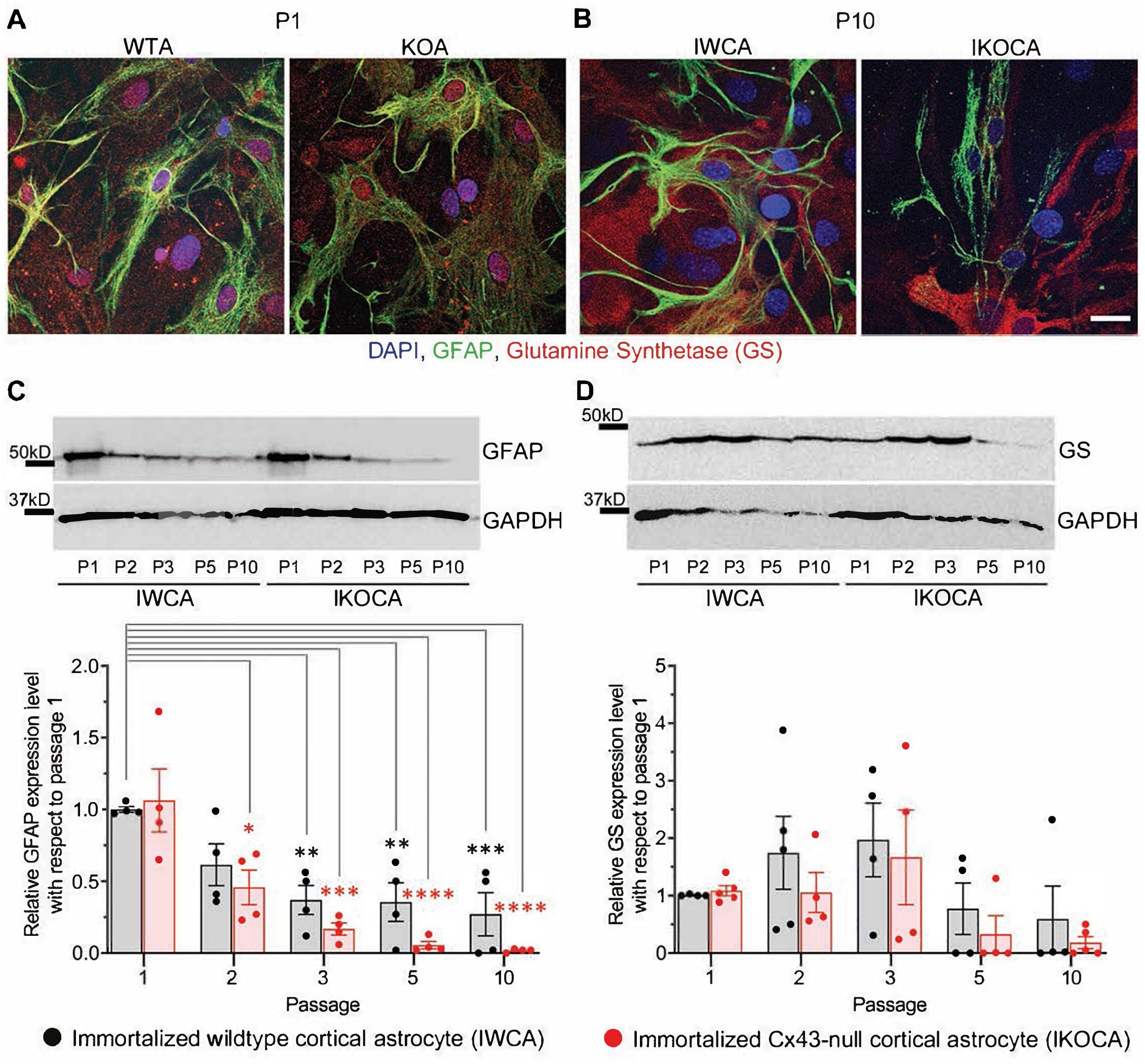
Expression level and localization of GFAP and GS in IWCA and IKOCA during the first ten passages. **A, B.** GFAP immunostaining declined with time in culture for both IWCA and IKOCA. GS was highly expressed in both IWCA and IKOCA cultures at all passages but declined at passage 5 in both genotypes. **C.** GFAP protein levels also declined rapidly with progressive passage, with minimal levels of about 27% in IWCA and about 2% in IKOCA cultures. **D.** GS protein levels increased over time in culture, being somewhat variable at all time points.

### Functional assessments of Cx43 expression in the immortalized cortical astrocyte cell lines

Our primary goal with the immortalization procedure was to obtain both normal astrocytes and astrocytes lacking expression of the major gap junction protein Cx43, which would provide an ideal tool to assess the role of Cx43 in multiple aspects of astrocyte behavior. To assure that functional Cx43 expression persisted in the immortalized astrocytes over multiple passages, we measured intercellular transfer of the small fluorescent dyes Lucifer Yellow (LY; MW 444 Da) and calcein (MW 622 Da). To quantify strength of coupling with LY, we injected individual cells in confluent culture dishes. As shown in Fig. 4A, at passage 9-10 LY transfer was common between IWCA, spreading from the injected cell to 13.7 ± 2.8% of its neighboring cells (Fig. 4C) in a total of 535 ± 100 cells per field. LY transfer in IKOCA cultures was virtually absent (Fig. 4B), spreading from the injected cell to only 1.0 ± 0.1% of its neighboring cells (Fig. 4C) in a total of 102 ± 27 cells per field. Dye transfer was also evaluated using the so-called “parachute assay”. In this assay, dissociated donor cells of passage 7-10 loaded with both the GJ permeable dye calcein and the lipophilic, GJ impermeable membrane dye DiI, were added dropwise onto the monolayer of recipient cells at passage 7-10. Coupling was observed at 1-2 hours after adding the donor cells. As shown in Figs. 4D and G, each donor IWCA cell recruited an average of neighboring 24.36 ± 8.95 cells in IWCA recipient cultures, confirming the findings from the LY injection assay demonstrating that IWCA are functionally coupled. Heterocellular coupling between IWCA and IKOCA was not observed (Figs. 4E, G), and the Cx43 deficient IKOCAs did not display homocellular coupling (Figs. 4F, G).

**Figure 4.**
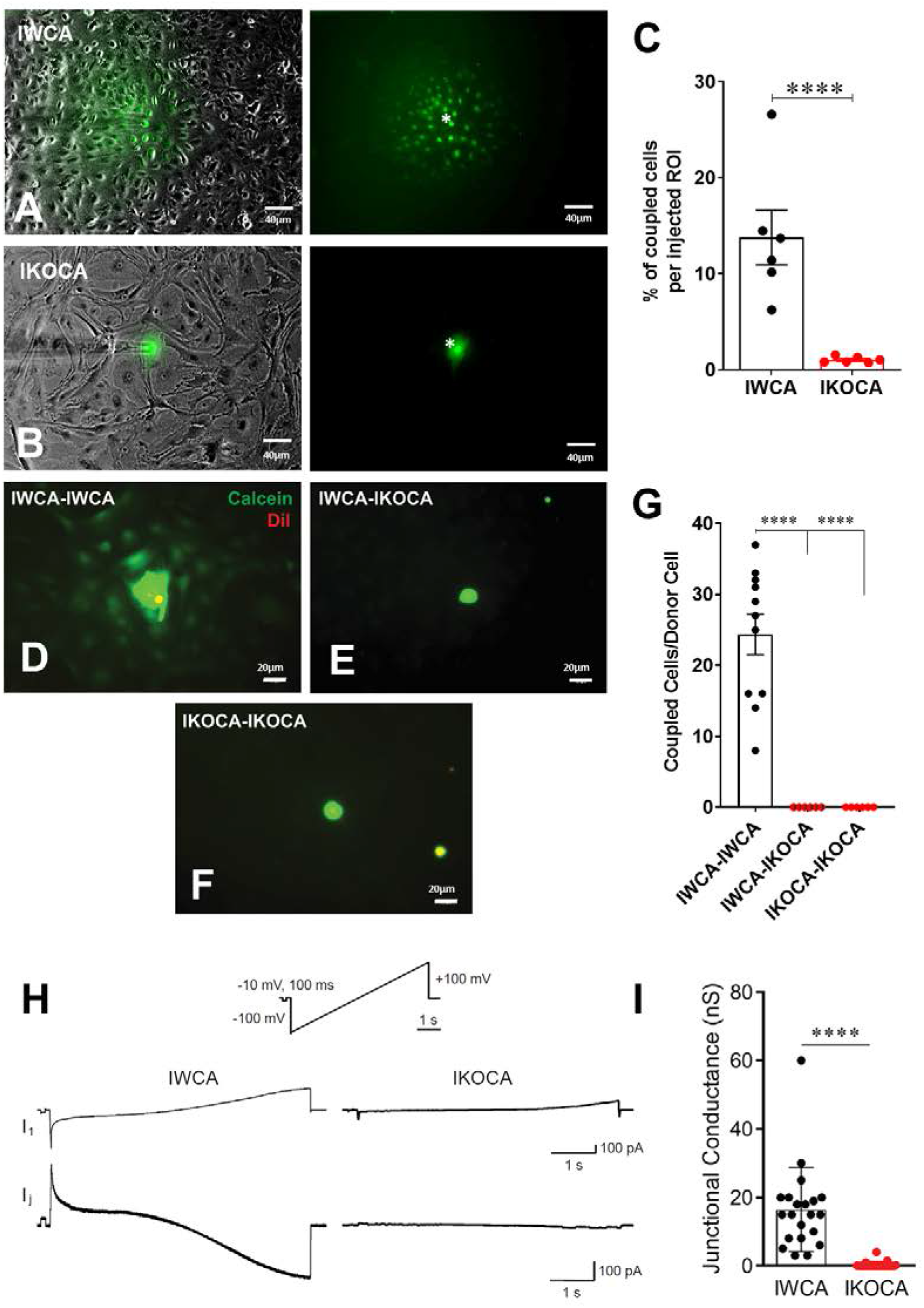
Assessment of functional gap junction coupling in immortalized WT (IWCA) and Cx43-null (IKOCA) cortical astrocyte cultures. **A.** Lucifer Yellow (LY) dye coupling between IWCA observed at passage 10. Left, bright phase contrast image of the field of view, with superimposed fluorescence. Right; fluorescence image with asterisk indicating the cell that was micro-injected with LY; note gap junction-mediated dye spread detected in all immediately adjacent neighbors. **B.** LY dye coupling between IKOCA observed at passage 10. Left, bright phase plus fluorescence showing field of view for injection. Right: In IKOCA cultures, the dye did not spread from the injected cell into the neighbor adjacent cells. **C.** Quantification of LY dye spread in IWCA and in IKOCA cultures. Bar histograms correspond to mean±SEM of 6 injections per genotype performed in two independent cultures per genotype at passage 10 (**** p<0.001). **D, E and F.** Parachute assay: **D.** from IWCA donor to recipient IWCA cells (IWCA-IWCA); **E.** from IWCA donor to recipient IKOCA cells (IWCA-IKOCA), and **F.** from IKOCA donor to recipient IKOCA cells. Donor cells were identified by double calcein-DiI labelling. Note extensive spread in IWCA-IWCA homotypic donor-recipient cells in D, and total lack of spread in the heterotypic IWCA-IKOCA and homotypic IKOCA-IKOCA in E and F, respectively. **G.** Quantification of calcein dye spread in IWCA-IWCA, in IWCA-IKOCA and in IKOCA-IKOCA donor-recipient cultures. Coupling was assessed as number of coupled cells per donor cell. **H.** Junctional currents measured using dual whole cell voltage clamp between IWCA and IKOCA cell pairs. Slow voltage ramp (inset, −100 mV to +100 mV over 5 sec) was delivered to cell 1, while recording currents in cell 1 (I1, upper traces) and cell 2 (current recorded in cell 2 while delivering voltage to cell 1 is equivalent, but opposite in polarity, to junctional current and is labeled Ij). Junctional current from IWCA pairs (lower left trace) are large and show nonlinearity with respect to voltage that is consistent with voltage dependence of Cx43 channels. Junctional current from IKOCA pair is quite small, indicating that cells are not highly coupled. **I.** Quantification of junctional conductance (gj = Ij/V) in IWCA and IKOCA cell pairs. Note significantly stronger gj in WCA compared to IKCA (17.35+3.2 nS, N=17 vs 0.36+0.18 nS, N=23; p<0.0001).

Electrophysiological recording using whole cell voltage clamp methods provides the most sensitive and quantitative method to measure GJ mediated coupling, and the biophysical properties thus revealed can indicate which connexins form the channels (9). Paired whole cell recordings from IWCA (Passage >10) in response to long voltages pulses showed junctional currents that were constant over time when voltage of either polarity was low, but current declined at voltage pulses higher than about ±40 mV (Fig. 4H). This weak but apparent voltage sensitivity of junctional conductance is similar to that measured in cell lines transfected with Cx43 (13). In IKOCAs (Passage >10), fewer cell pairs were coupled and junctional conductance was much lower (Fig. 4I). Calculated junctional conductance in IWCA was 17.5+3.2 nS in 17 cell pairs, compared to 0.36+.18 nS in 23 IKOCA cell pairs, a substantial and significant difference (p <0.0001).

Findings from the dye coupling and electrophysiological studies demonstrate the preserved ability of IWCA to form functional Cx43 GJ channels. In the IKOCA, the inability to form functional Cx43 GJs is expected to be solely attributable to the lack of Cx43 expression. To demonstrate that the cellular machinery needed to assemble and form Cx43 GJs is also preserved in IKOCA, we transfected these cells with GFP-tagged Cx43 and compared GJ plaque formation with that of IWCA. As shown in Fig 5A, gap junctions outline the margins of IWCAs but are totally absent in non-transfected IKOCAs (Fig. 5D). In IWCA, the endogenous Cx43 GJ plaques appear to consist of somewhat discrete smaller plaques when viewed at higher magnification (Fig. 5B and 5C). Similarly, Cx43 plaques in IKOCAs transfected with GFP-tagged Cx43 appear as oblate domains containing circular subunits (Fig. 5E and 5F). The areas of the cell in Fig. 5E with dimmer but detectable GFP signal represent non-junctional sfGFP-Cx43 that can be recognized as localized mostly to the plasma membrane when viewed in 3D at high magnification using a confocal microscope. Overall increased expression of Cx43 and larger gap junction plaque structures represent differences noticeable for exogenously expressed connexins and endogenously expressed Cx43 found in IWCA astrocytes. These results indicate that immortalization of Cx43-null astrocytes did not impair the ability of these cells to form GJs, and that the organization of GJ plaques in IKOCA transfected with GFP-tagged Cx43 resemble the GJ plaques endogenously expressed between IWCA. Most importantly, this finding indicates that the IKOCA cell line may be a useful substrate in which Cx43 gap junctions and their variants can be expressed in an astrocyte background to emulate a normal cellular context in order to investigate effects of Cx43 variant-expression on cell physiology and morphology absent the confound of endogenous Cx43 expression.

**Figure 5.**
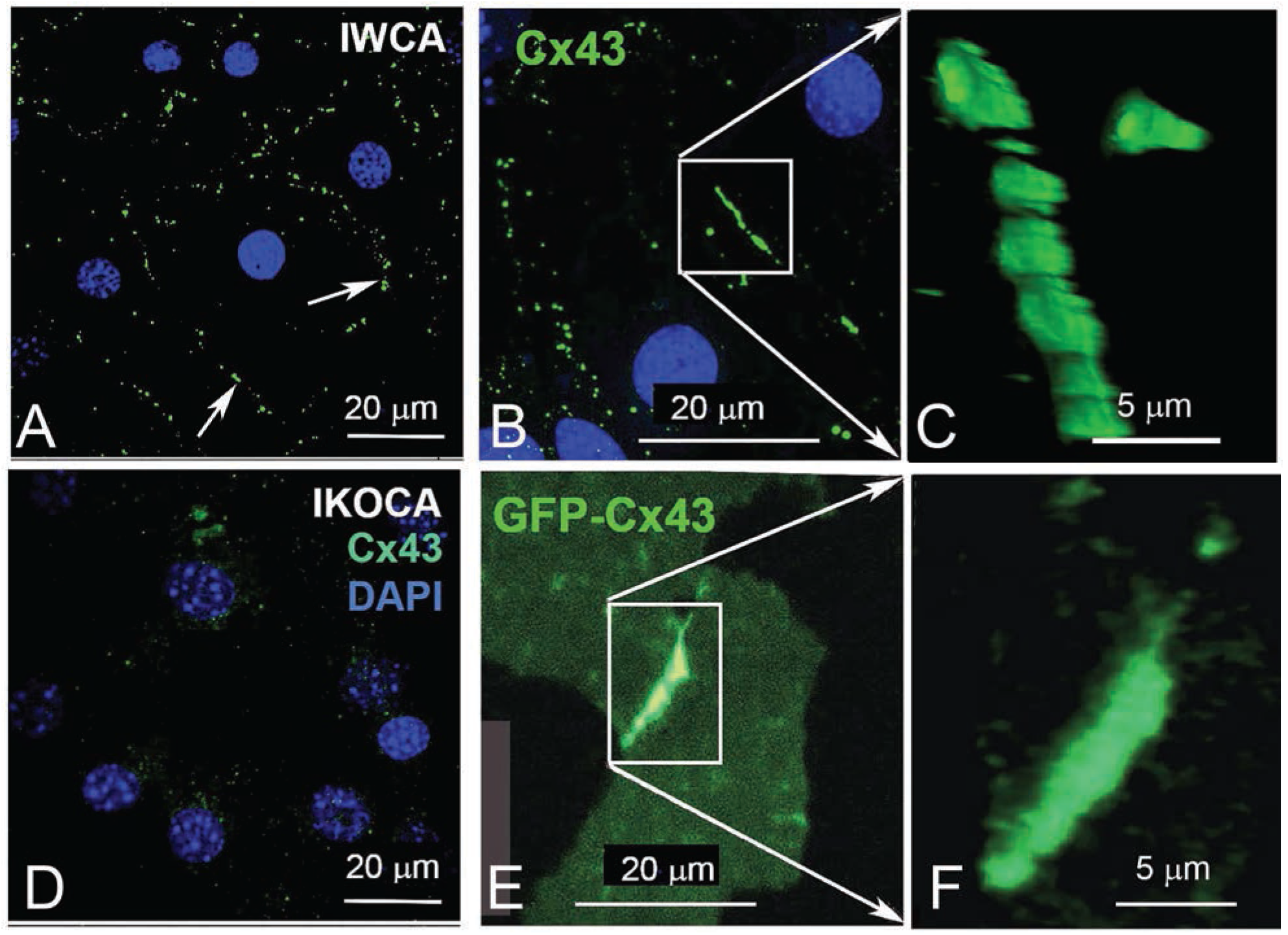
Comparison of endogenous Cx43 gap junction plaques in immortalized WT cortical astrocytes (IWCA) with GJ plaques formed by GFP-tagged Cx43 in immortalized Cx43-null cortical astrocytes (IKOCA). **A.** Gap junction plaques (arrows) outline individual IWCAs (identified by DAPI-stained nuclei). **B** and **C.** At higher magnification, these junctional domains appear somewhat oblate, and are formed by closely packed plaques. **D.** In IKOCAs, endogenous Cx43 junctions are absent. **E.** Following transfection with GFP-tagged Cx43, plaques are induced between cell IKOCA pairs. **F.** At higher magnification, these appositional domains appear oblate, composed of a series of smaller plaques.

### IKOCA cells allow super-resolution imaging of gap junction plaque morphology and interactions with other organelles

The locations of EBFP2-Cx43 (immunostained green in Figure 6) vesicles (green puncta) and gap junction plaques connecting two IKOCA cells were imaged using a Zeiss 980 Airyscan2 microscope to examine of gap junction plaque morphological features such as discontinuities and small neighboring gap junction plaques (Zoomed inset in Figure 6A). Note the small bright puncta near the edge of the gap junction plaque shown in figure 6A, inset in which morphology reminiscent of budding endocytic gap junctions observed previously via electron microscopy and super-resolution light microscopy by others in transformed cell lines (14). The interactions between gap junctions and mitochondria and with other organelles modify intercellular communication (15, 16). The re-expression of Cx43 in some IKOCA cells within a culture as shown Figure 6 allows comparison of the effect of Cx43 expression in IKOCA cells on mitochondrial location and morphology. Here we show post-fixation immunofluorescence staining of all mitochondria. Note the conspicuous localization of mitochondria to areas of the EBFP2-Cx43 plaque (Fig 6C shows a reflexive Cx43 GJ plaque, white arrow) that is partially undergoing endocytosis with small mitochondria potentially located within the large endocytic portions of the gap junction plaque (GJ endosomes also known as connexosomes). Connexosome-associated mitochondria are indicated by yellow arrows in Figure 6D. Areas where the astrocyte membrane is folded but not occupied by gap junction plaque is indicated by the faint but enhanced green signal next to the small white arrowheads in Figure 6C. These figures highlight the utility of IKOCA and IWCA astrocytes for high resolution imaging.

**Figure 6.**
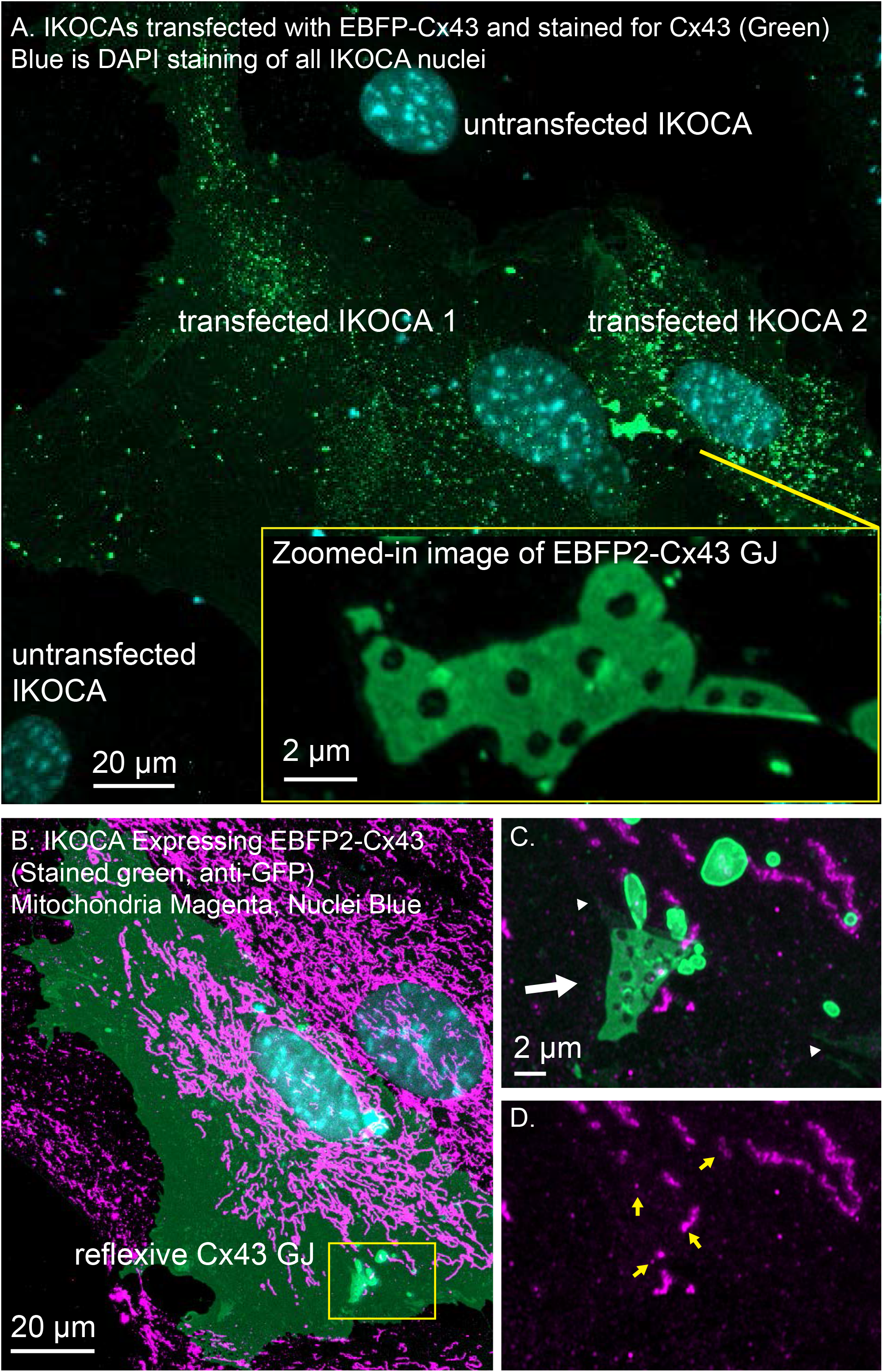
IWCA and IKOCA cells are useful to study gap junction morphology and interactions with other proteins with super-resolution microscopy. **A.** IKOCA cells transfected to re-express Cx43 with a fluorescent protein tag (green, immunolabeling with an antibody against Cx43) form gap junctions when two transfected cells are in contact. As indicated by blue DAPI staining, there are untransfected cells in contact with the two transfected IKOCA cells in the center of the field of the image in A that do not express Cx43 and do not form GJs. Super-resolution imaging allows insights into structural features of the gap junction plaque and endocytosed GJ plaque as shown in the zoomed inset. **B.** Re-expression of fluorescent protein tagged Cx43 in a subset of IKOCA cells with subsequent immunostaining for secondary cellular features present in all IKOCA cells (e.g. mitochondria) allows investigation of the effects of Cx43 expression in astrocyte-like cells within the same cell culture sample-facilitating side-by-side comparisons and visualization of interactions between Cx43 and other organelles with super-resolution microscopy. **C.** Zoomed in view of the location where a Cx43 gap junction plaque structure (green, indicated by white arrow) appears to interact with small mitochondria (indicated by TOMM20 staining, magenta). This gap junction plaque represents a reflexive gap junction plaque that has formed where the plasma membrane of the same astrocyte overlaps to allow gap junction formation. **D.** View of the same region as in “C.” with the Cx43 channel removed to allow visualization of the TOMM20 staining within endocytic Cx43 connexosomes (yellow arrows)-raising the possibility that mitochondrial transfer may occur via Cx43 endocytosis.

### IKOCA cells allow optical experimentation on gap junctions and interacting proteins

Three-dimensional FRAP such as that shown in Figure 7A allows improved assessment of intra-GJ plaque protein mobility since the entire effective (for this time-scale) fluorescence pool available to contribute to recovery can be detected in 3D FRAP. The motion of other gap junction features such as discontinuities that form when part of the gap junction is endocytosed can be easily studied in immortalized astrocytes using a confocal microscope. The arrangement of Cx43 is much more stable within the gap junction plaque structure than other proteins such as Cx30 and Cx26, as reported previously in transformed cell lines (Stout et. al. 2015). The thin, flat morphology of the cultured IKOCA and IWCA cells leads them to produce gap junction plaques that often align nearly parallel with the growth substrate (coverslip). Such characteristics make IKOCA and IWCA cells ideal for live 3D microscopy and super-resolution imaging of gap junctions and intracellular organelles. Ability to entirely replace the absent Cx43 expression in IKOCA cells with fluorescent protein tagged Cx43 allows co-expression of other proteins of interest tagged with fluorescent proteins in compatible color channels. Figure 7,B shows an orthogonal maximum intensity projection of a 3D time-lapse image of an pair of IKOCA cells joined by a blue-fluorescent (pseudo colored magenta) EBFP2-Cx43 gap junction plaque that is partially undergoing endocytosis. The cells were co-transfected with msfGFP-ATG9A (green). ATG9A was previously shown to interact with the endocytosis process of Cx43 and to be colocalized to the plasma membrane (Bejarano et. al. 2014) but the apparent transfer of small vesicle-like structures positive for ATG9A to portions of the Cx43 gap junction plaque at the point of endocytosis initiation (Figure 7B, zoomed inset) was not previously reported and would be very difficult to observe in astrocyte like cells without access to IKOCA cells.

**Figure 7.**
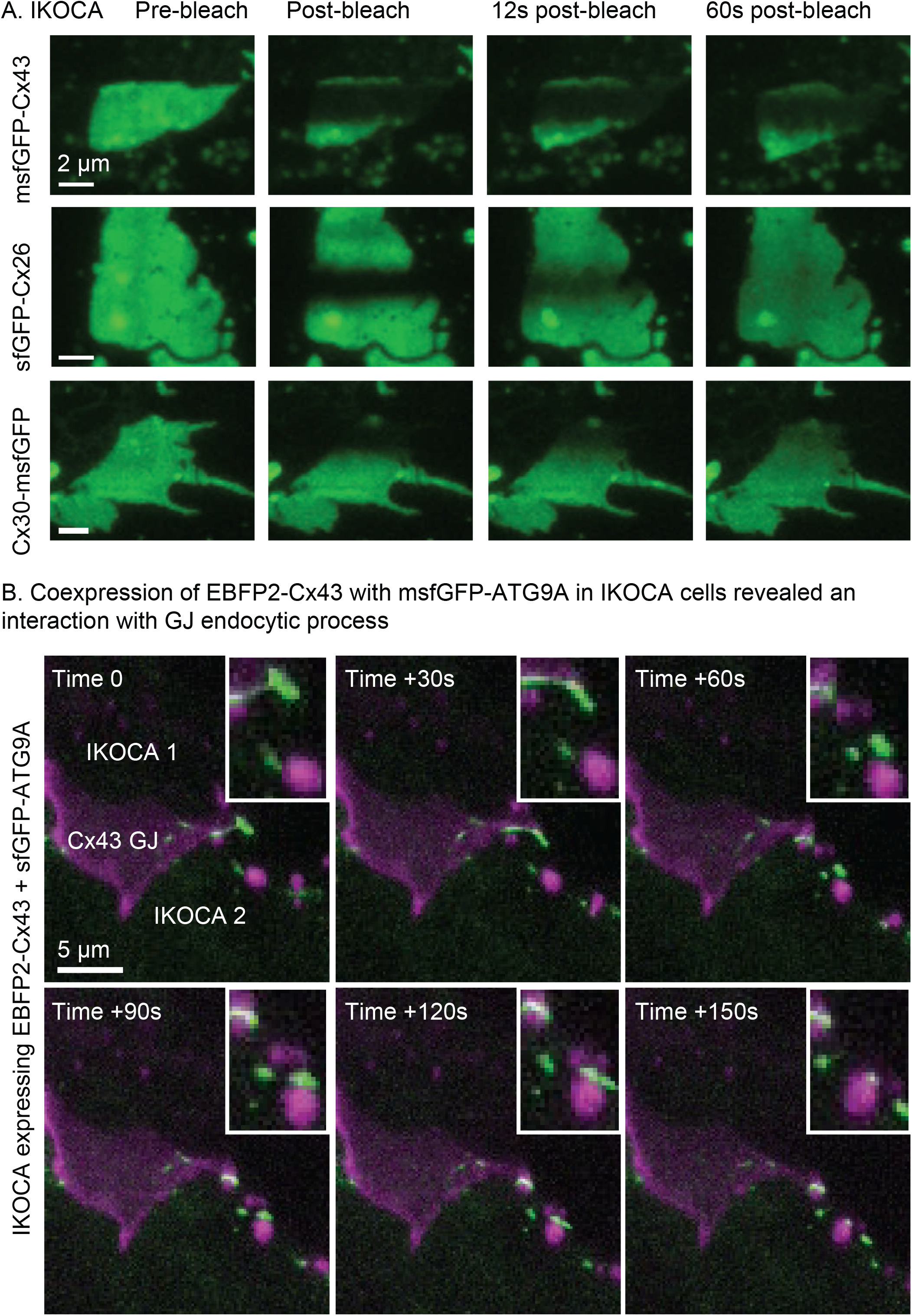
Expression of modified Cx43 transgenes in IKOCA cells can be visualized live without the confounding effects of endogenous Cx43 expression and allows testing the effects of expression of Cx43 mutants and other treatments on IWCAs. **A.** Expression of sfGFP-Cx43 and other fluorescent protein-tagged connexins allows 3D time lapse FRAP experiments to examine gap junction plaque dynamics in astrocyte-like cells. The thin morphology of astrocytes facilitates 3D, time lapse FRAP since gap junction plaques often form in a nearly parallel configuration with the imaging focal plane. For the example images shown, the full vertical width of the gap junction was imaged using 11 confocal planes with each stack acquired at 3 second intervals. The resulting Z-stacks were collapsed to a series of maximum projection reconstructions to visualize the fluorescence recovery for the entire plaque structure over time. **B.** Expression of EBFP2-Cx43 with green fluorescent protein tags and sensors allows live, 4D examination of interactions between Cx43 and other cell components such as msfGFP-ATG9A displayed as maximum-projections as a time-lapse montage. Please note that EBFP2-Cx43 gap junction plaque is shown in magenta with the msfGFP-ATG9A autophagy protein shown in green. The zoomed inset shows the detail of the dynamic interaction between transgene-expressed Cx43 and ATG9A in IKOCA astrocytes.

### Barrier formation in endothelial monolayer is enhanced by co-culture with IWCAs

To evaluate whether the immortalized astrocyte cell lines retain their ability to functionally interact with endothelial cells, as observed at the level of the blood-brain barrier, we conducted studies with IWCA co-cultured with endothelial bEnd.3 cells. Barrier formation was evaluated based on development of Trans Endothelial Electrical Resistance (TEER) that was monitored for several days using the CellZscope apparatus. As seen in Figure 8, the TEER of cultures of bEnd.3 endothelial cells with and without IWCAs increased with time in culture. However, TEER of cultures of bEnd.3 cells with IWCA was significantly higher (p-value<0.05) than either bEnd.3 alone or together with IKOCA since the beginning of the measurement and persisting until day 7. By contrast, IKOCA-bEnd.3 co-cultures formed a much leakier endothelium. At 7 days, the resistance of the IWCA-bEnd.3 co-cultures was about two and a half times as high as that measured with bEnd.3 cells alone [23.17 ± 2.60 Ohm-cm^2^ for co-cultures compared to 9.34 ± 0.86 Ohm-cm^2^ for bEnd.3 cells alone; (p=0.0003)], whereas the Cx43 null astrocytes formed a barrier that was much lower (12.56 ± 1.13 Ohm-cm^2^). These IWCA co-cultures TEER values are similar to those previously reported by others at 5 days from co-cultures of bEnd.3 cells with primary rat astrocytes (17) using the Endohm chamber connected to an EVOM resistance meter (18).

**Figure 8.**
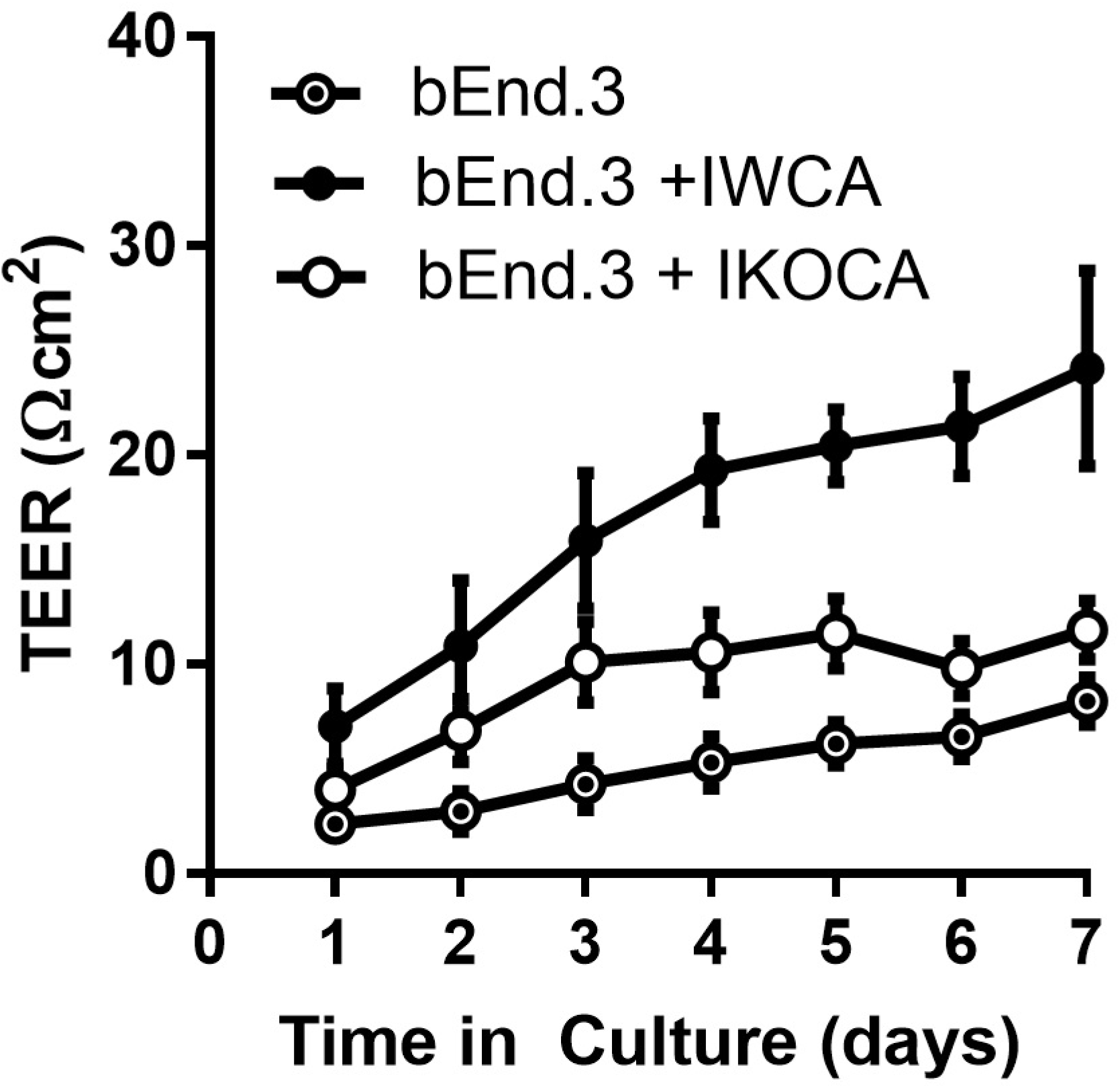
Transwell resistance (TEER) measured in bEnd.3 endothelial cells alone and in co-culture with immortalized WT (IWCA) and with Cx43-null (IKOCA) cortical astrocytes. Increase in electrical resistance of the transwell cultures of bEnd.3 cells alone and in co-culture with IWCA and IKOCA measured using the CellZscope. N=11-17 separate experiments. Since day 1, the TEER (Ohm cm^2^) of co-cultures with IWCA was significantly higher than bEnd.3 cells alone (7.31 ± 1.13 vs 1.67 ± 0.28, p-value = 0.00024) and IKOCA with bEnd.3 (7.31 ± 1.13 vs 4.33 ± 0.63, p-value = 0.033). On day 7 the difference persisted when compared with bEnd.3 cells alone (23.17 ± 2.60 vs 9.34± 0.86, p-value = 0.00027) and with IKOCA with bEnd.3 (23.17 ± 2.60 vs 12.56 ± 1.13, p-value = 0.0022).

## Discussion

The immortalized cell lines that we have developed express biomarkers consistent with that of the astrocyte phenotype. In the IWCA lines, Cx43 mRNA level was rather constant during ten passages, although protein level declined slightly at passage 10. Cx43 was absent in the immortalized null line. Cx30 was absent in the immortalized cells of both genotypes; this is likely consistent with its very low expression in neonatal astrocytes, where it only becomes detectable in older brains and in astrocytes cultured for long periods (19). Cx26, by contrast, was expressed at moderately high levels in early cultures and declined with passage. Likewise, the water channel that characterizes the astrocyte endfoot, AQP4, was expressed at highest level in early passages and its mRNA and protein declined to low but still detectable levels by passage 10. The mRNAs and protein levels of four biomarkers by which the astrocyte phenotype is usually assessed, ALDH1, GS, GLT-1 and GFAP, displayed both similarities and differences. ALDH1 and GS mRNAs were relatively constant during successive passages, whereas protein levels were constant or, in the case of GS, actually increased until passage 10. Expression of glutamate transporter GLT-1 and GFAP mRNA declined with passage; GFAP protein level declined but was still detectable at passage 10. Immunostaining at corresponding times substantiated the Western blots and also indicated that expression was in appropriate compartments within the cells.

Gap junction mediated intercellular communication was maintained in the IWCA line and was very low in the IKOCA line. With respect to transfer of small molecular weight dyes, both intracellularly injected Lucifer Yellow and ester loaded calcein were highly permeable to junctions between cells. Electrophysiological measurements of electrical coupling found junctional conductance averaging 17 nS between IWCA cell pairs, which is similar to the value of 13 nS that we have previously reported in primary cultures of these cells (6). Voltage sensitivity of junctional conductance, a distinctive biophysical property of gap junctions formed of each connexin, was similar to that reported for Cx43 (6).

The development of methods that permitted mammalian cells to be maintained in culture for prolonged periods of time was a major accomplishment that pioneered the fields of cell biology and cellular neuroscience. Cell types from some tissues continue to be much more difficult to extract and maintain in culture than oncogenically transformed cell lines. Most differentiated primary cells extracted from tissue undergo a limited number of cell divisions before becoming senescent, necessitating continuous cell harvesting to maintain a supply of material dissociated from tissues for study. The most commonly used transformed cell lines have contributed greatly to cell biological research but have a number of well-known drawbacks including major changes to cell phenotypes, signaling pathways and characteristics of intercellular interactions. It has thus been desirable to obtain cell lines that divide continuously and which also express phenotypes resembling the cell type of interest.

Conventional methods to obtain immortalized cell lines have involved the isolation of cells from tumors [see(20), examples of lines initially generated (earlier than 1960) being a mouse lung cell line (L1), the highly discussed and widely used human cervical cell line (HeLa; (21)) and Chinese hamster ovary cells (CHO)], but thousands of other cell lines have since been derived from tumors and characterized to some extent or another. It is now known that these cell lines arose from oncogene-expressing cells. HeLa cells are infected with human papillomavirus (HPV), which synthesizes proteins that inactivate the tumor suppressor molecule p53 and others (22).

Studies of mechanisms responsible for tumorigenesis have exploited SV40, a polyomavirus that was discovered in a monkey cell line used to produce poliovirus vaccine. SV40 infects monkey cells, activating expression of early and late genes, and does not produce tumors in monkeys. In rodent cells, SV40 generates both large and small T-antigens to inhibit pRb and p53 as well as a protein phosphatase (PP2A) (23). In mice, SV40 virus infection leads to the induction of tumors and has been used to generate a number of widely used mouse cell lines (reviewed in (24)). In cell culture, SV40 infection leads to cell transformation, generally assayed as loss of contact inhibition of growth, ability to grow in soft agar and decreased requirement of serum for survival. Most cell lines in current use were generated by this method and thus exhibit the transformed phenotype. For example, the American Tissue Cell Collection repository contains seven astrocyte cell lines. One of these DI TNC-1, obtained by GFAP-targeted oncogene delivery to neonatal rat diencephalon astrocytes (25), has low levels of endogenous aquaporins and has thus been a useful model to study exogenous constructs expressed in an astrocyte-similar transformed cell line [e.g. (26)]. Products are even commercially available tailored specifically to such studies (https://altogen.com/product/di-tnc1-transfection-reagent-rat-brain-astrocytes/).

A more recent alternative for the immortalization of primary cells involves the incorporation of hTERT, which preserves length of telomeres and thus endows cell with properties of self-renewal. hTERT introduction has an advantage over the use of viral protein for immortalization in that it does not involve the inactivation of tumor suppressor gene, thereby bypassing the transformed phenotype (27). Thus, hTERT immortalization results in cells that typically more closely resemble the primary cells than do virus-transformed cells, although both immortalization methods escape the limited number of cell cycles before senescence and may not totally mimic the properties of the cells from which they are derived.

Most studies using cell lines to evaluate consequences of gap junction expression on astrocyte phenotype have used C6 glioma cells (ATCC #CCL-107). This cell line was derived from rat brain tumors generated by nitrosomethylurea exposure and was initially characterized as rich in S100 protein and spindle-shaped (28). The C6 cell line is widely used in studies of glioblastoma invasion, as it forms robust tumors *in vivo*. It has been described as containing two populations of cells, one having markers of immature oligodendrocytes and another being a mixture of oligodendrocyte and immature astrocytes (29). Notably, GFAP expression is absent and vimentin expression is variable in C6 glioma cells (30). With regard to gap junction research, C6 glioma cells were found to express less Cx43 than astrocytes, and are regarded as being severely communication deficient (31). In a number of studies, overexpression of Cx43 in these cells has been reported to decrease tumor growth rate *in vivo* and cell proliferation in culture (see (32)). However, C6 cells form functional gap junctions with astrocytes (33), and numerous studies have more recently indicated that gap junction communication though Cx43 channels plays a major role in glioblastoma invasion (see (34)).

Few studies directly related to the issue of tumorigenesis have been conducted on astrocytes derived from Cx43 deficient astrocytes. However, it is clear that manipulation of Cx43 expression in C6 cells may have very different consequences than in astrocytes. For example, in a study comparing gene expression levels in WT C6 cells with three Cx43 transfected clones, only seven distinct genes were found to be up or downregulated (35). In transcriptome-wide studies comparing astrocytes from WT mice with either astrocytes from Cx43-null mice (36) or following Cx43 knockdown with siRNA(37), hundreds of genes were up or downregulated. Gene regulation in Cx43-null and Cx43 knockdown astrocytes was to some extent similar (37), but none of the genes identified in the transfected C6 cells was regulated in either of the Cx43-null or Cx43 knockdown astrocytes.

The cell lines developed here exhibit characteristics of primary astrocytes and are expected to offer new opportunities for the evaluation of the roles of astrocyte gap junctions than cells derived from glioblastomas or other tumors. For example, future studies examining binding partners and their affinity for astrocyte proteins (38) are likely to be more similar to those that occur *in vivo* when examined in a more natural context (although the presence of low Cx26 expression in IKOCA cultures should be considered). And the degree to which properties of astrocytes depend on Cx43 expression and identification of molecular domains responsible should be readily accessible in astrocyte cell lines derived from sibling WT and Cx43-null mice.

## Abbreviations

GJ: Gap Junction
Cx43: Connexin43
Cx30: Connexin30
Cx26: Connexin26
AQP4: Aquaporin4
GS: glutamine synthetase
GFAP: Glial Acidic Fibrillary Protein
GLT-1 (EAAT2): Glutamate transporter 1 (Excitatory Amino Acid Transporter 2)
qRT-PCR: quantitative real-time polymerase chain reaction
hTERT: human telomerase reverse transcriptase
ALDH1: aldehyde dehydrogenase 1
bEnd.3 cells: mouse brain microvascular endothelial cell line ATCC® CRL-2299™
IWCA: immortalized wildtype cortical astrocytes
IKOCA: immortalized Cx43-null cortical astrocytes
sfGFP: superfolder green fluorescent protein
EBFP2: enhanced blue fluorescent protein 2

## Other acknowledgements

This study was conducted as a joint project by our laboratories, and we acknowledge this collaboration by alphabetization of the authorship. Some data was acquired with equipment of the Analytical Imaging Facility at Albert Einstein College of Medicine which is supported in part by an NCI Cancer Center Support Grant P30CA013330. Some data was acquired or processed with equipment of the Center for Biomedical Innovation at the New York Institute of Technology, College of Osteopathic Medicine. The astrocyte cell lines described in this paper are available to the scientific community without charge, subject to standard MTAs with the New York Institute of Technology College of Osteopathic Medicine.

